# Neuroanatomical and functional dissociations between variably present anterior lateral prefrontal sulci

**DOI:** 10.1101/2023.05.25.542301

**Authors:** Ethan H. Willbrand, Silvia A. Bunge, Kevin S. Weiner

**Author notes:** Corresponding author Kevin S. Weiner. Co-senior authorship (Silvia A. Bunge, Kevin S. Weiner).

## Abstract

The lateral prefrontal cortex (LPFC) is an evolutionarily expanded region in humans that is critical for numerous complex functions, many of which are largely hominoid-specific. While recent work shows that the presence or absence of specific sulci in anterior LPFC is associated with cognitive performance across age groups, it is unknown whether the presence of these structures relates to individual differences in the functional organization of LPFC. To fill this gap in knowledge, we leveraged multimodal neuroimaging data from 72 young adult humans aged 22-36 and show that dorsal and ventral components of the paraintermediate frontal sulcus (pimfs) present distinct morphological (surface area), architectural (thickness and myelination), and functional (resting-state connectivity networks) properties. We further contextualize the pimfs components within classic and modern cortical parcellations. Taken together, the dorsal and ventral pimfs components mark transitions in anatomy and function in LPFC, across metrics and parcellations. These results emphasize that the pimfs is a critical structure to consider when examining individual differences in the anatomical and functional organization of LPFC and highlight the importance of considering individual anatomy when investigating structural and functional features of the cortex.

## Introduction

A main goal in cognitive and systems neuroscience is to precisely understand how the human cerebral cortex is organized morphologically, anatomically, and functionally. Of particular interest are association cortices, which have expanded the most throughout evolution and present anatomical and functional features that are cognitively relevant – some of which are unique to humans. For example, classic and ongoing work shows that the lateral prefrontal cortex (LPFC) displays a complex structural and functional organization that supports numerous complex cognitive abilities (Badre & D’Esposito, 2009; Demirtaş et al., 2019; Levy & Goldman-Rakic, 2000; Nee & D’Esposito, 2016; Petrides, 2005; Rosenkilde, 1979; Stuss & Knight, 2013). A growing body of recent work demonstrates the utility of studying small, shallow, and variable sulci (often referred to as tertiary sulci; Armstrong et al., 1995; Chi et al., 1977; Sanides, 1964; Welker, 1990) for understanding the anatomical and functional organization of association cortices, including LPFC (Amiez et al., 2013, 2021; Amiez & Petrides, 2014; Y. Li et al., 2015; Lopez-Persem et al., 2019; Miller, D’Esposito, et al., 2021; Miller, Voorhies, et al., 2021; Sanides, 1964; Troiani et al., 2016, 2020; Weiner, 2019; Willbrand, Parker, et al., 2022). Intriguingly, some tertiary sulci are present in every brain, while others are not (Amiez et al., 2019; Hathaway et al., 2023; Malikovic et al., 2012; Miller et al., 2020; Miller, Voorhies, et al., 2021; Nakamura et al., 2020; Paus et al., 1996; Petrides, 2019; Vallejo-Azar et al., 2022; Willbrand, Parker, et al., 2022; Willbrand, Voorhies, et al., 2022). In the present study, we focus on the morphological, architectural, and functional features of variably present sulci in anterior LPFC—the dorsal (pimfs-d) and ventral (pimfs-v) components of the paraintermediate frontal sulcus (pimfs), respectively. We do so for four main reasons.

First, the anterior and posterior LPFC differ based on incidence rates of the small, shallow, and variable tertiary sulci located within them. Across age groups, posterior LPFC contains three tertiary sulci that are present in all participants (Miller, Voorhies, et al., 2021; Voorhies et al., 2021; Yao et al., 2022). By contrast, in anterior LPFC, a given hemisphere can have (i) a pimfs-d and pimfs-v, (ii) a pimfs-d, but not a pimfs-v (or vice versa), or (iii) neither component (Willbrand, Jackson, et al., 2023; Willbrand, Voorhies, et al., 2022). Second, the sulcal depth of a subset of these posterior and anterior LPFC sulci are related to cognitive performance (Voorhies et al., 2021; Yao et al., 2022). Third, two separate studies in pediatric and adult cohorts show that the presence or absence of the pimfs is related to reasoning performance (Willbrand, Jackson, et al., 2023; Willbrand, Voorhies, et al., 2022). Fourth, while our prior work indicated that the three posterior LPFC sulci are anatomically distinct structures that co-localize with distinct functional networks (Miller, Voorhies, et al., 2021), the anatomical and functional distinctiveness and relevance of the pimfs components have yet to be investigated.

Therefore, to fill this gap in knowledge, we tested whether the two pimfs components are functionally and/or anatomically dissociable. To do so, we applied classic multimodal criteria (Felleman & Van Essen, 1991; Kaas, 1997; Van Essen, 2003) to 249 pimfs labels from 72 participants from the Human Connectome Project (HCP; 144 hemispheres; 50% female, aged 22-36) via a three-pronged approach. First, we extracted and compared the morphological (depth, surface area) features of the pimfs components. Second, we did the same for architectural (gray matter thickness, myelination) features of the pimfs. Third, we created functional connectivity profiles for each pimfs component using functional network parcellations of the human cerebral cortex unique to each HCP participant that was created blind to cortical folding and our sulcal definitions (Kong et al., 2019). Finally, we contextualized the alignment of our individual-level pimfs labels with several widely used group-level modern and classic parcellations of the human cerebral cortex spanning multiple cortical features.

## Materials and Methods

### Multimodal HCP dataset

Data for the young adult human cohort analyzed in the present study were taken from the Human Connectome Project (HCP) database: ConnectomeDB (db.humanconnectome.org). Here, as in several prior studies (Miller, Voorhies, et al., 2021; Willbrand, Jackson, et al., 2023; Willbrand, Parker, et al., 2022), we used a randomly selected subset of 72 participants (50% female, aged between 22 and 36 years old), given the time-intensive process of individual sulcal labeling. Additionally, previous work examining structural-functional correspondences in individual hemispheres shows that this sample size is large enough to encapsulate individual differences and detect reliable effects in individual hemispheres (e.g., as few as 20 hemispheres is typically considered a sufficient sample size; (Amiez et al., 2006; Amunts et al., 2020; Amunts & Zilles, 2015; Lopez-Persem et al., 2019; Zlatkina et al., 2016). HCP consortium data were previously acquired using protocols approved by the Washington University Institutional Review Board and informed consent was obtained from all participants.

Anatomical T1-weighted (T1-w) MRI scans (0.7 mm voxel resolution) were obtained in native space from the HCP database (db.humanconnectome.org), along with outputs from the HCP modified FreeSurfer pipeline (v5.3.0; (Dale et al., 1999; Fischl, Sereno, & Dale, 1999; Fischl, Sereno, Tootell, et al., 1999; Glasser et al., 2013). Additional details on image acquisition parameters and image processing can be found in the previously published work by Glasser and colleagues (Glasser et al., 2013). Maps of the ratio of T1-w and T2-w scans, which is a measure of tissue contrast enhancement related to myelin content, were downloaded as part of the HCP ‘Structural Extended’ release. All subsequent sulcal labeling and extraction of anatomical metrics were calculated on the cortical surface reconstructions of individual participants generated through the HCP’s custom-modified version of the FreeSurfer pipeline (Dale et al., 1999; Fischl, Sereno, & Dale, 1999; Fischl, Sereno, Tootell, et al., 1999; Glasser et al., 2013).

### Anatomical analyses

#### Manual sulcal labeling

LPFC sulci were manually defined within each individual hemisphere using tksurfer, as in prior work (Miller, Voorhies, et al., 2021; Voorhies et al., 2021; Willbrand, Ferrer, et al., 2023; Willbrand, Jackson, et al., 2023; Willbrand, Voorhies, et al., 2022; Yao et al., 2022). Manual lines were drawn on the inflated cortical surface to define sulci based on the most recent schematics of pimfs and sulcal patterning in LPFC by Petrides (Petrides, 2019), as well as by the pial and smoothwm surfaces of each individual (Miller, Voorhies, et al., 2021). In some cases, the precise start- or end-point of a sulcus can be difficult to determine on a surface (Borne et al., 2020). Thus, using the inflated, pial, and smoothwm surfaces to inform our labeling allowed us to form a consensus across surfaces and clearly determine each sulcal boundary. The location of pimfs components was confirmed by trained independent raters and finalized by a neuroanatomist (K.S.W.).

In the present study, we restricted our analyses to the anterior MFG (aMFG; **Figure 1**), as the anatomical and functional properties of the tertiary sulci in posterior MFG (pMFG) have already been assessed (Miller, Voorhies, et al., 2021). Although this project focused primarily on the pimfs and three immediately surrounding sulci [i.e., the horizontal component of the intermediate middle frontal sulcus (imfs-h), ventral component of the intermediate middle frontal sulcus (imfs-v), and inferior frontal sulcus (ifs)], the manual identification of the other 19 LPFC sulci (2,985 sulcal definitions across all 72 participants) was required to ensure the most accurate definition of all sulci. For in-depth descriptions of all LPFC sulci, see (Miller, D’Esposito, et al., 2021; Miller, Voorhies, et al., 2021; Petrides, 2019; Voorhies et al., 2021; Willbrand, Ferrer, et al., 2023; Willbrand, Jackson, et al., 2023; Yao et al., 2022). In each hemisphere, we first labeled the surrounding primary (ifs) and secondary sulci (imfs-h and imfs-v) so that we could use them as landmarks to identify the pimfs (**Figure 1**). As described in prior work (Willbrand, Jackson, et al., 2023; Willbrand, Voorhies, et al., 2022), the dorsal and ventral components of the pimfs (pimfs-d and pimfs-v) were generally defined using the following two-fold criterion: i) the sulci ventrolateral to the imfs-h and imfs-v, respectively, and ii) superior and/or anterior to the mid-anterior portion of the ifs (**Figure 1**).

**Figure 1.**
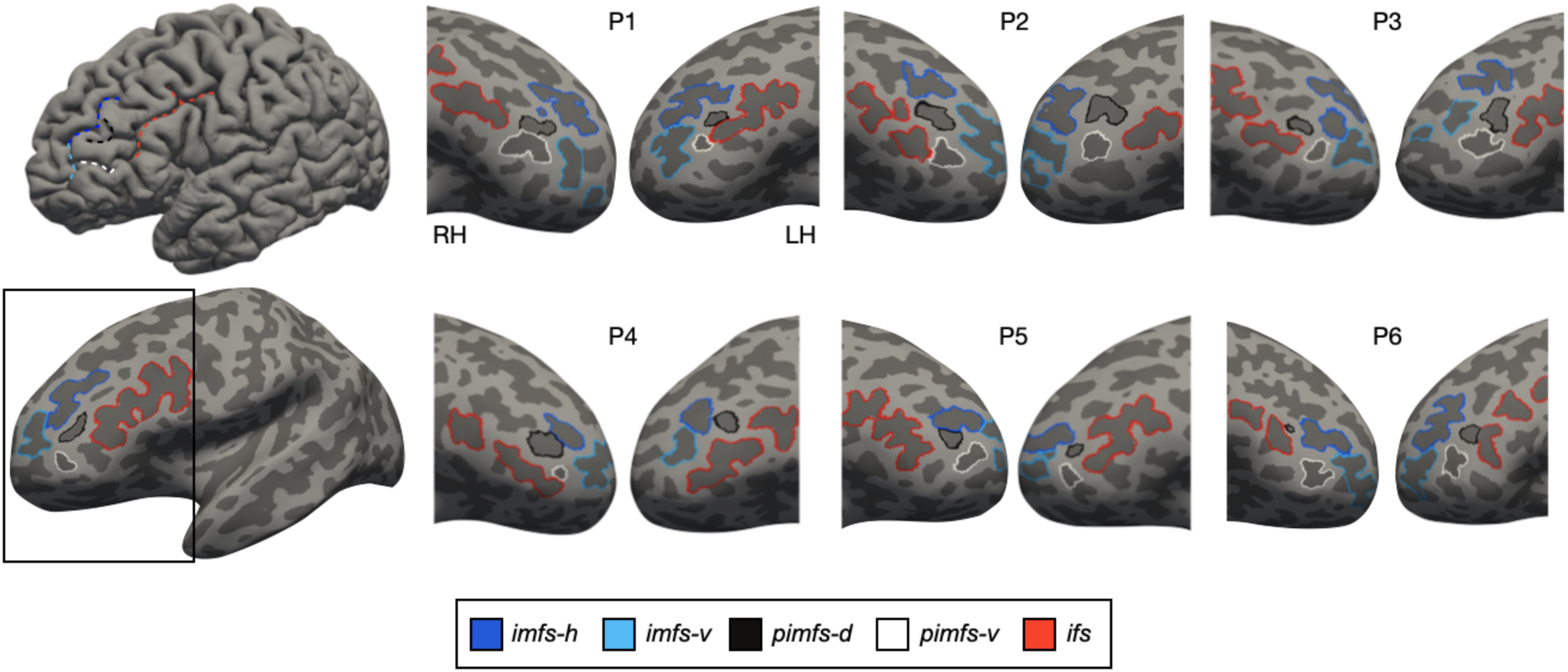
Components of the paraintermediate frontal sulcus are often, but not always, identifiable within individual hemispheres. Left: pial (top) and inflated (bottom) left hemisphere (dark gray: sulci; light gray: gyri) from an example participant with the two components of the paraintermediate frontal sulcus (dorsal: pimfs-d; ventral: pimfs-v) defined, as well as three prominent surrounding sulci: i) horizontal component of the intermediate frontal sulcus (imfs-h), ii) ventral component of the intermediate frontal sulcus (imfs-v), and inferior frontal sulcus (ifs). Sulci are colored according to the key below. The black box around the inflated surface focuses on the lateral prefrontal cortex (LPFC). Right: Additional left (LH) and right (RH) inflated cortical surfaces of six individual participants focused on the LPFC. While there can be 0, 1, or 2 pimfs components in a given hemisphere, we primarily show hemispheres containing 2 components (with the exception of P4). 57% of individuals had both components in both hemispheres.

#### Quantifying and comparing the morphology and architecture of the paraintermediate frontal sulcus components

Morphologically, we compared the depth and surface area of the pimfs components, as these are two of the primary morphological features used to define and characterize sulci (Armstrong et al., 1995; Chi et al., 1977; X. Li et al., 2022; Lopez-Persem et al., 2019; Madan, 2019; Miller, D’Esposito, et al., 2021; Miller et al., 2020; Miller, Voorhies, et al., 2021; Natu et al., 2021; Petrides, 2019; Sanides, 1964; Voorhies et al., 2021; Weiner, 2019; Weiner et al., 2014, 2018; Welker, 1990; Willbrand, Ferrer, et al., 2023; Willbrand, Parker, et al., 2022; Willbrand, Voorhies, et al., 2022; Yao et al., 2022). We expected that the pimfs components would be shallower and smaller than the three more prominent sulci surrounding them, based on our prior work on the three pMFG tertiary sulci in young adults (Miller, Voorhies, et al., 2021) and for the pimfs in children and adolescents (Voorhies et al., 2021). Indeed, this is what we found (**Figure A.1**).

Sulcal depth and surface area were measured following the same procedures as in our prior work (Voorhies et al., 2021; Yao et al., 2022). Mean sulcal depth values (in standard FreeSurfer units) were computed in native space from the .sulc file generated in FreeSurfer (Dale et al., 1999; Fischl, Sereno, & Dale, 1999; Fischl, Sereno, Tootell, et al., 1999) with custom Python code (leveraging functions from the nilearn and nibabel packages) developed in our prior work (Voorhies et al., 2021). Briefly, depth values are calculated based on how far removed a vertex is from what is referred to as a “mid-surface,” which is determined computationally such that the mean of the displacements around this “mid-surface” is zero. Thus, generally, gyri have negative values, while sulci have positive values. Given the shallowness and variability in the depth of tertiary sulci (Miller, Voorhies, et al., 2021; Voorhies et al., 2021; Yao et al., 2022), some mean depth values extend below zero. We emphasize that this just reflects the metric implemented in FreeSurfer. Each depth value was also normalized by the deepest point in the given hemisphere. Surface area (in square millimeters) was generated for each sulcus through the mris_anatomical_stats function in FreeSurfer (Fischl & Dale, 2000). Surface area was normalized by hemispheric surface area as in our prior work (Hathaway et al., 2023; Willbrand, Ferrer, et al., 2023; Willbrand, Voorhies, et al., 2022).

Architecturally, we compared cortical thickness and myelination (**Figure 2A**), as in our prior work (Miller, Voorhies, et al., 2021; Voorhies et al., 2021; Willbrand, Parker, et al., 2022). Mean gray matter cortical thickness (mm) was extracted from each sulcus using the mris_anatomical_stats function in FreeSurfer (Fischl & Dale, 2000). To quantify myelin content, we used an in vivo proxy of myelination: the T1-w/T2-w maps for each individual hemisphere (Glasser & Van Essen, 2011; Shams et al., 2019). To generate the T1-w/T2-w maps, two T1-w and T2-w structural MR scans from each participant were registered together and averaged as part of the HCP processing pipeline (Glasser et al., 2013). The averaging helps to reduce motion-related effects or blurring. Additionally, the T1-w/T2-w images were bias-corrected for distortion effects using field maps, as described by Glasser and colleagues (Glasser et al., 2013). We then extracted the average T1-w/T2-w ratio values across each vertex for each sulcus using custom Python code, leveraging functions from the nilearn and nibabel packages (Miller, Voorhies, et al., 2021).

**Figure 2.**
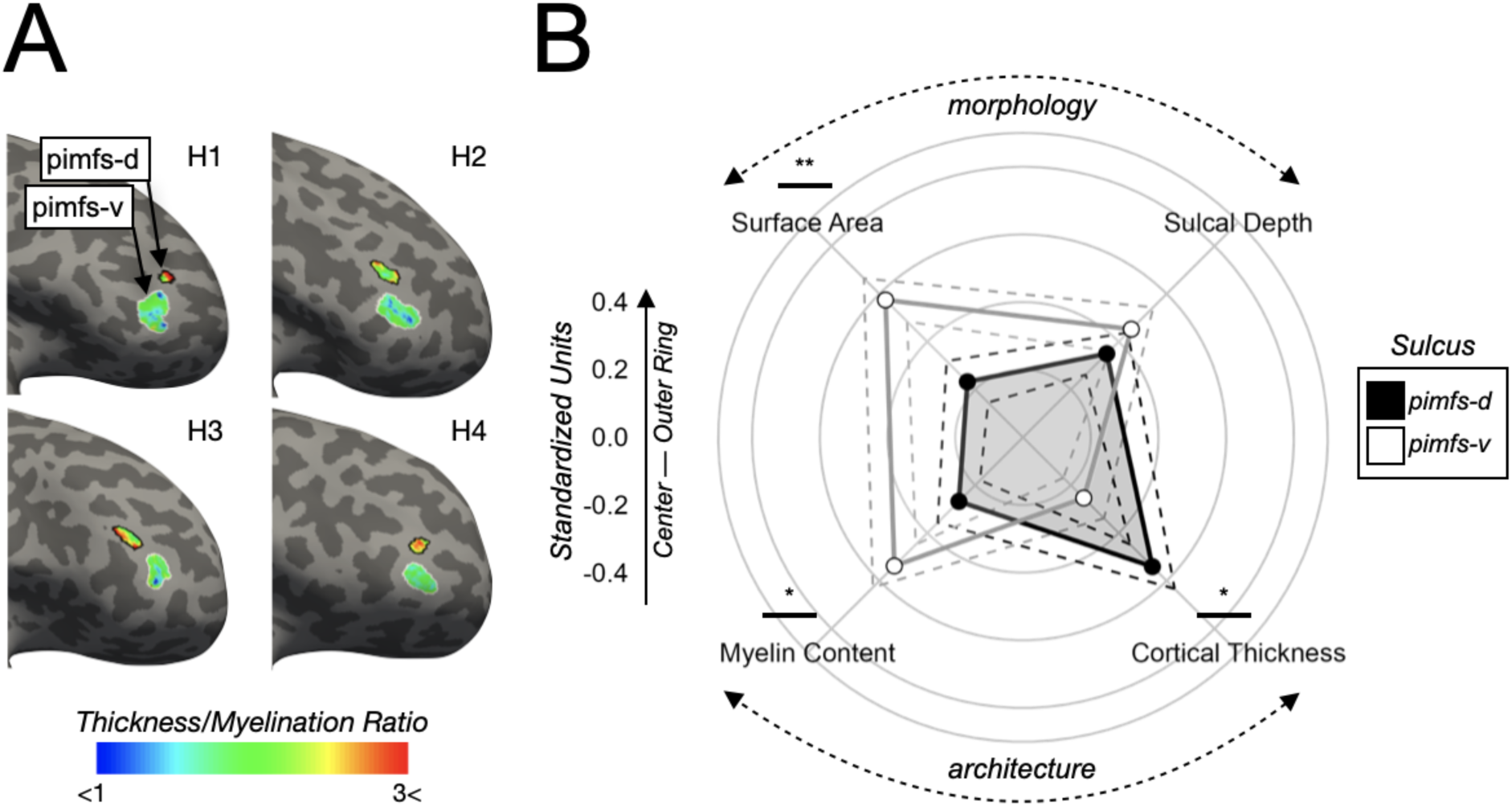
Pimfs-d is 15.73% smaller, 2.48% cortically thicker, and 0.81% less myelinated on average than pimfs-v. **A.** Four example inflated hemispheres (labeled H1, H2, etc; two left and two right; all oriented as right hemispheres) displaying the thickness/myelination ratio (heatmap; see bottom color bar) within each pimfs component (pimfs-d: black outline; pimfs-v: white outline). Surfaces are focused on LPFC as in Figure 1. Note that these example hemispheres display the effects shown in B: the pimfs-d is smaller, as well as thicker and less myelinated (as shown by the higher thickness/myelination ratio) than the pimfs-v. **B.** Polar plot showing the mean morphological (top) and architectural (bottom) values for the pimfs components (averaged across hemisphere; see **Figure A.2** for these values split by hemisphere). Solid lines and dots represent the means. Dashed lines represent ± standard error. Lines and dots are colored by sulcal component (pimfs-d: black, pimfs-v: white/gray). Each concentric circle corresponds to the units shown to the left, which are standardized to allow for these metrics to be plotted together. Line and asterisks above each of the metric labels indicate the post hoc pairwise comparisons on the sulcus × metric interaction (* *p* < .05, ** *p* < .01).

To assess whether these four metrics differed between the pimfs components, we ran a linear mixed effects model (LME) with the following predictors: sulcal component (pimfs-d and pimfs-v) × metric (surface area, depth, cortical thickness, and myelination) × hemisphere (left and right). Sulcal component, metric, and hemisphere were treated as fixed effects. Metric was nested within sulcus, which was nested within hemisphere, which was nested within subject. An ANOVA F-test was subsequently conducted, from which results were reported. We also assessed whether the presence/absence of the pimfs-d impacted these features of the pimfs-v, and vice versa, by running an LME for each component, exchanging the predictor “sulcal component” for “number of components” (one, two).

### Functional analyses

To assess whether the pimfs components are functionally distinct, we implemented a three-pronged approach leveraging data spanning the individual (Kong et al., 2019), meta-analysis (Yeo et al., 2015), and group levels (Fan et al., 2016; Foit et al., 2022; Glasser et al., 2016; Scholtens et al., 2018; Van Essen, 2005), which we now discuss in turn.

#### Individual level: Comparing connectivity of the paraintermediate frontal sulcus components from resting-state functional connectivity network parcellations

To determine whether the pimfs components are functionally distinct, we generated functional connectivity profiles, or “connectivity fingerprints”, using a recently developed analytic approach (Miller, Voorhies, et al., 2021; Willbrand, Parker, et al., 2022). First, we used resting-state network parcellations for each individual participant from Kong and colleagues (Kong et al., 2019), who previously generated individual network definitions by applying a hierarchical Bayesian network algorithm to produce maps for each of the 17 networks in individual HCP participants. These data were calculated in the template HCP fs_LR 32k space. Importantly, this parcellation was conducted blind to cortical folding (and therefore, our sulcal definitions). Next, we resampled the network profiles for each participant onto the fsaverage cortical surface, and then to each native surface using CBIG tools (https://github.com/ThomasYeoLab/CBIG).

We then calculated, for each hemisphere and participant, the spatial overlap between a sulcus and each of the (i) eight main networks comprising the parcellation (Auditory, Control, Default, Dorsal Attention, Somatomotor, Temporal-Parietal, Ventral Attention, Visual) and (ii) 17 individual resting-state networks (i.e., considering sub-networks: Auditory, Control A, Control B, Control C, Default A, Default B, Default C, Dorsal Attention A, Dorsal Attention B, Somatomotor A, Somatomotor B, Temporal-Parietal, Ventral Attention A, Ventral Attention B, Visual A, Visual B, Visual C). To quantify the overlap between a sulcus and each of the networks, we computed Dice coefficients:

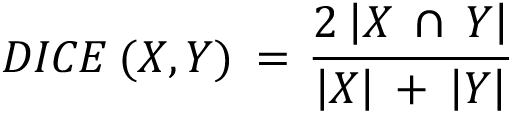

where *X* and *Y* are the sulcus and network, | | represents the number of elements in each set, and ∩ represents the intersection of two sets. Fourth, we ran two LMEs [predictors: sulcal component (pimfs-d and pimfs-v) × network (8 or 17 networks) × hemisphere (left and right)] to determine whether the network profiles (i.e., the Dice coefficient overlap with each network) of the pimfs-d and pimfs-v were differentiable from one another. In the first LME, we compared the profiles using the general eight networks to assess broad correspondences. Next, we compared the connectivity fingerprints of the pimfs components with all 17 networks to determine which sub-networks were driving the effect in the first model. In this second model, sulcal component, network, and hemisphere were treated as fixed effects. Network was nested within sulcal component, which was nested within hemisphere, which was in turn nested within subject. ANOVA F-tests were applied to each model.

To quantify variability and individual differences in the connectivity fingerprints of each pimfs component, we calculated the Wasserstein metric (Earth Mover’s Distance) between the resting-state network overlap values for each unique pair of participants, such that a larger distance indicates decreased similarity. We then applied the non-parametric Wilcoxon signed-rank test to the pimfs Wasserstein metric data to assess whether the pimfs components differed in terms of inter-individual variability of the pattern of network overlap.

#### Meta-analysis: Cognitive component modeling of the paraintermediate frontal sulcus components

To further assess the functional dissociability of the pimfs components, we quantified the overlap between each sulcal component and meta-analytic fMRI data at the group level from 10,449 neuroimaging experiments, which yielded 14 probabilistic “cognitive component” maps (Yeo et al., 2015). The cognitive component model from Yeo and colleagues (Yeo et al., 2015) links patterns of brain activity to behavioral tasks via latent components representing putative functional subsystems. Each cognitive component map, which was calculated on the fsaverage surface, provides the probability that a given voxel will be activated by one of the 14 components (Yeo et al., 2015).

To relate the sulcal components of each individual to the vertex-wise maps of the 14 cognitive components (Yeo et al., 2015), we first aligned each sulcal label to the fsaverage surface, using the mri_label2label FreeSurfer function. We then used a Bayesian method of expectation maximization to determine the combination of cognitive components that best fit each sulcal component for each participant in each hemisphere. This resulted in a set of probabilities of overlap between the pimfs-d and pimfs-v and each of the 14 cognitive components. We previously validated this approach by showing that the somatomotor components of the cognitive component map aligned strongly with the central sulcus (Miller, Voorhies, et al., 2021). Moreover, we showed that it was possible to detect functional differences between neighboring tertiary sulci: specifically, between the three pmfs components in the pMFG (Miller, Voorhies, et al., 2021). Therefore, in the present study, we tested whether the pimfs components were distinguishable based on these cognitive component loadings, via an LME [predictors: sulcal component (pimfs-d and pimfs-v) × cognitive component (14 components) × hemisphere (left and right)]. Sulcal component, cognitive component, and hemisphere were treated as fixed effects. Cognitive component was nested within sulcal component, which was nested within hemisphere, which was nested within subject. ANOVA F-tests were applied to each model. These results are discussed in **Figure A.5**.

#### Group level: Comparing co-localization of the paraintermediate frontal sulcus components with classic and modern group-level parcellations of the cerebral cortex

Finally, we sought to situate the pimfs components with respect to modern and classic cortical parcellations. In the main text we highlight two parcellations: the group-level HCP 180-region multimodal parcellation (HCP-MMP), derived from topography, architecture, function, and connectivity (Glasser et al., 2016), as well as Brodmann’s cytoarchitectonic parcellation (Brodmann, 1909) mapped onto the fsavarage surface (i.e., the PALS B12 Brodmann atlas; (Van Essen, 2005). We specifically focused on the HCP-MMP because it is based on multiple anatomical and functional metrics, was derived from the sample used in the present study, and has been highly influential since its release (Glasser et al., 2016). We also focused on Brodmann’s cytoarchitectonic parcellation because it is foundational to the field of brain mapping, having been used to identify the location of different functional areas in thousands of studies (Zilles, 2018).

We adopted a similar procedure to the one used for the individually derived parcellations and meta-analysis cognitive components described above. First, we resampled the pimfs components of each participant to the common fsaverage surface, which the HCP-MMP and Brodmann parcellations were also mapped onto (Glasser et al., 2016; Van Essen, 2005). Second, for each participant and hemisphere, we calculated the Dice coefficient to measure the overlap between each sulcal component and the group-level parcellations in the Glasser and Brodmann atlases that comprise LPFC: specifically, eight HCP-MMP regions (IFS-p, IFS-a, p9-46v, 46, 9-46d, a9-46v, p47r, a47r; (Glasser et al., 2016) and six Brodmann Areas (BAs; 45, 46, 47, 9, 10, 11; (Brodmann, 1909). Third, we ran an LME [predictors: sulcal component (pimfs-d and pimfs-v) × ROI × hemisphere (left and right)] to determine if the the pimfs-d and pimfs-v were differentiable from one another based on each parcellation. ROI was nested within sulcal component, which was nested within hemisphere, which was nested within subject. ANOVA F-tests were applied to each model.

We then repeated this pipeline with three additional cortical parcellations, to expand upon the two focused on in the main text. First, we used the modern, functional connectivity-based Brainnetome atlas (specifically nine LPFC regions: IFJ, A8vl, A9/46d, A9/46v, IFS, A46, A45r, A10l, A12/47l; (Fan et al., 2016). Second, we used the classic Von Economo and Koskinas (von Economo & Koskinas, 1925) cytoarchitecture parcellation—specifically five LPFC regions: FC, FD, FDdelta, FDT, FF—which was recently projected to the fsaverage surface by Scholtens et al. (Scholtens et al., 2018). Third, we used the classic myeloarchitecture parcellation of the Vogt-Vogt school (Vogt & Vogt, 1919) (specifically seven LPFC regions: 48, 49, 52, 53, 54, 58, 59), which was also recently projected to the fsaverage surface by Foit et al. (Foit et al., 2022). The same format of LME was applied in these cases as well.

### Statistics

All statistical tests were implemented in R (v4.0.1). LMEs were implemented with the lme function from nlme R package. ANOVA F-tests were run with the anova function from the stats R package. Effect sizes for the ANOVAs are reported with the partial eta-squared (η2) metric. Relevant post hoc pairwise comparisons on ANOVA effects were computed with the emmeans and contrast functions from the emmeans R package (*p*-values adjusted with Tukey’s method). The effect size for post hoc pairwise comparisons is reported with the Cohen’s d (*d*) metric. Wasserstein distance was calculated with the wasserstein1d function from the transport R package. The Wilcoxon test was implemented with the wilcox.test function from the stats R package. If an effect or interaction with a factor (such as hemisphere) is not explicitly reported, it is not significant.

## Results

### When present, the dorsal and ventral components of the pimfs differ morphologically and architecturally

As described in the **Materials and Methods** and in our prior work (Willbrand, Jackson, et al., 2023; Willbrand, Voorhies, et al., 2022), the pimfs components are two variable sulci in the aMFG, identified based on their proximity to the more prominent and superior imfs (**Figure 1**). The dorsal pimfs is inferior to the horizontal imfs, whereas the ventral pimfs is inferior to the ventral imfs (**Figure 1**). Both sulci are superior and anterior to the ifs (**Figure 1**). The pimfs is also variably present across the 72 young adult participants in this sample (see example hemispheres in **Figure 1**): in a given hemisphere, individuals may have 2, 1, or 0 components. In this sample, the pimfs-d was present in 89% of the left and 88% of the right hemispheres, whereas the pimfs-v was present in 81% of the left and 89% of the right hemispheres (Willbrand, Jackson, et al., 2023). With regard to the number of components present, both pimfs-d and pimfs-v were present in in 72% of the left and 78% of the right hemispheres, a single one was present in 25% of the left and 21% of the right hemispheres, and neither was present in 3% of the left and 1% of right hemispheres.

After defining the pimfs components, we tested, based on four metrics, whether they differed morphologically and architecturally (**Materials and Methods**). Morphologically, we tested sulcal surface area (normalized to hemispheric surface area) and depth (normalized to maximal hemispheric depth), since these are two of the primary features used to describe sulci (e.g., (Armstrong et al., 1995; Chi et al., 1977; X. Li et al., 2022; Lopez-Persem et al., 2019; Madan, 2019; Miller, Voorhies, et al., 2021; Natu et al., 2021; Petrides, 2019; Sanides, 1964; Weiner, 2019; Welker, 1990). Architecturally, we assessed cortical thickness (in mm) and myelination (T1-w/T2-w ratio; (Glasser et al., 2013; Glasser & Van Essen, 2011); see **Figure 2A** for these values displayed on example hemispheres), as they are additional metrics commonly used to describe and compare sulci (e.g., (Alemán-Gómez et al., 2013; Ammons et al., 2021; Bertoux et al., 2019; Fornito et al., 2008; Miller et al., 2020; Miller, Voorhies, et al., 2021; Natu et al., 2019; Voorhies et al., 2021; Willbrand, Ferrer, et al., 2023; Willbrand, Parker, et al., 2022; Yao et al., 2022).

An LME [predictors: sulcal component (pimfs-d and pimfs-v) × metric (surface area, depth, cortical thickness, and myelination) × hemisphere (left and right)] revealed a sulcal component × metric interaction (F(3, 735) = 5.50, η2 = 0.02, *p* = .001). Post hoc pairwise comparisons revealed that (i) the pimfs-d was on average 15.73% smaller than the pimfs-v (*d* = 0.34, *p =* .005), (ii) there were no differences in sulcal depth (*d* = 0.10, *p* = .38), (iii) the pimfs-d was on average 2.48% cortically thicker than the pimfs-v on average (*d* = 0.26, *p* = .020), and (iv) the pimfs-d was on average 0.81% less myelinated than the pimfs-v (*d* = 0.30, *p* = .029; **Figure 2B**). Removing outliers did not meaningfully impact this interaction or the subsequent post hoc comparisons; in fact, doing so made some of the differences numerically stronger (interaction (F(3, 714) = 6.67, η2 = 0.03, *p* < .001), surface area (*d* = 0.38, *p* = .001; pimfs-d 16.36% smaller than pimfs-v on average), depth (*d* = 0.10, *p* = .32), cortical thickness (*d* = 0.35, *p* = .015; pimfs-d 2.92% thicker than pimfs-v on average), and myelination (*d* = 0.33, *p* = .010; pimfs-d 0.81% less myelinated than pimfs-v on average)). Further, the surface area, depth, cortical thickness, and myelination of the pimfs-d and pimfs-v did not differ based on the presence/absence of the other component (*p*s > .31). Altogether, these results indicate that the pimfs-d and pimfs-v are dissociable on the basis of morphology (surface area) and architecture (cortical thickness and myelination) at the individual level.

### When present, the ventral and dorsal components of the pimfs are functionally dissociable

Classic and recent work implicate the topography of tertiary sulci in the functional organization of association cortices (Amiez et al., 2013; Amiez & Petrides, 2014; Y. Li et al., 2015; Lopez-Persem et al., 2019; Miller, D’Esposito, et al., 2021; Miller, Voorhies, et al., 2021; Sanides, 1964; Troiani et al., 2016, 2020; Weiner, 2019; Willbrand, Parker, et al., 2022). Particularly relevant to the present study, our prior work indicated that the pMFG tertiary sulci were dissociable based on their relationship to fMRI connectivity networks (Miller, Voorhies, et al., 2021). Therefore, we sought to extend this assessment to the pimfs components in the aMFG.

To this end, we leveraged individual-level resting-state functional connectivity parcellations in the HCP sample (Kong et al., 2019). Importantly, these individual-level parcellations were developed without consideration for cortical folding (and therefore blind to our sulcal labels). For each pimfs component, we calculated the overlap with 8- and 17-functional network parcellations via the Dice coefficient (**Materials and Methods**). Akin to prior work on individual-level functional network variations (Gordon et al., 2017; Seitzman et al., 2019), this procedure generated a “connectivity fingerprint” for each pimfs component for each participant that is reflective of whole-brain connectivity patterns (for an example of this individual-level sulcal-network overlap see **Figure 3A**, see **Figures A.3 and A.4** for all individual connectivity fingerprints).

**Figure 3.**
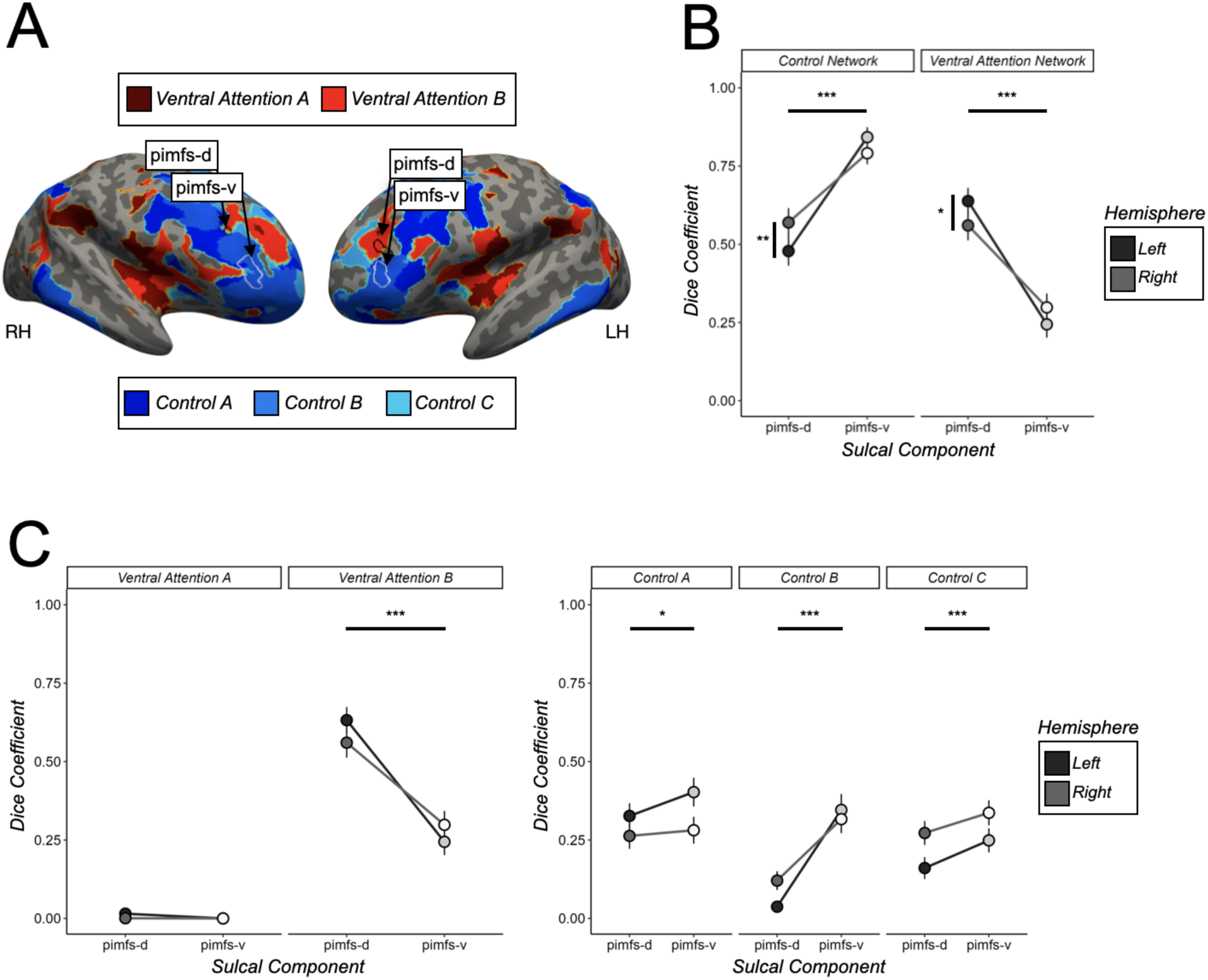
The pimfs components differ based on individually-derived functional connectivity fingerprints. **A.** Example left (LH) and right (RH) hemispheres displaying the relationship between the pimfs components (pimfs-d: black outline; pimfs-v: white outline) and the Ventral Attention/Salience networks (red areas), as well as the Control networks (blue areas) as defined by Kong et al., 2019. We only visualize these two broad networks/five sub-networks, as they are the only ones prominently overlapping with the pimfs. **B.** Dice coefficients are plotted as a function of sulcal component (x-axis; pimfs-d: black, pimfs-v: white), broad networks (facets), and hemisphere (left hemisphere: darker shades; right hemisphere: lighter shades). Large dots and error bars represent mean ± standard error. Horizontal lines and asterisks indicate the significance of the post hoc pairwise comparisons stemming from the sulcus × network interaction on Dice coefficient overlap (* *p* < .05, ** *p* < .01, *** *p* < .001). Vertical lines and asterisks indicate the significance level of the post hoc pairwise comparisons stemming from the sulcus × network × hemisphere interaction. **C.** Same as B, but for sub-networks: the Ventral Attention/Salience and Control sub-networks. Lines and asterisks indicate the significance level of the post hoc pairwise comparisons stemming from the sulcus × network interaction on Dice coefficient overlap. Although there was a sulcus × network × hemisphere for the broad networks, this interaction was not significant with the sub-networks (*p* = .19).

We first assessed the relationship between the pimfs components and eight broad functional connectivity networks identified by Kong and colleagues (Kong et al., 2019); Auditory, Control, Default, Dorsal Attention, Somatomotor, Temporal-Parietal, Ventral Attention/Salience, Visual). An LME [predictors: sulcal component (pimfs-d and pimfs-v) × network (8 networks) × hemisphere (left and right)] revealed a sulcal component × network interaction (F(7, 1659) = 55.09, η2 = 0.19, *p* < .001). Post hoc pairwise comparisons revealed a double dissociation: pimfs-d overlapped more with the Ventral Attention/Salience network (*d* = 0.95, *p* < .001; pimfs-d: mean ± se = 0.60 ± 0.03, pimfs-v: mean ± se = 0.27 ± 0.03; **Figure 3B**), whereas pimfs-v overlapped more with the Control network (*d* = 0.92, *p* < .001; pimfs-d: mean ± se = 0.52 ± 0.03, pimfs-v: mean ± se = 0.82 ± 0.02; **Figure 3B**). There was also a sulcal component × network interaction × hemisphere interaction (F(7, 1659) = 2.78, η2 = 0.01, *p* = .007), such that the pimfs-d overlapped more with the Control network in the right hemisphere (*d* = 0.25, *p* = .004) and with the Ventral Attention/Salience network in the left hemisphere (*d* = 0.22, *p* = .015), thereby indicating that the dissociation was stronger in the left hemisphere (**Figure 3B**). Additionally, the pattern of overlap between a given component and the networks did not differ based on whether or not the other component was present (*p*s > .40).

We then assessed the degree of overlap of the pimfs components with the sub-networks of these aforementioned networks (Control A, Control B, Control C, Ventral Attention/Salience A, Ventral Attention/Salience B; **Figure 3A**). Once again, an LME [predictors: sulcal component (pimfs-d and pimfs-v) × network (17 networks) × hemisphere (left and right)] revealed a sulcal component × network interaction (F(16, 3792) = 26.93, η2 = 0.10, *p* < .001). Post hoc pairwise comparisons revealed that the pimfs-d overlapped more with Ventral Attention/Salience B sub-network (*d* = 0.94, *p* < .001; **Figure 3C**), while the pimfs-v overlapped more with the three Control sub-networks: Control A (*d* = 0.13, *p* = .022), Control B (*d* = 0.86, *p* < .001), and Control C (*d* = 0.27, *p* < .001; **Figure 3C**). It is worth noting that the overlap of pimfs-d with the broad Ventral Attention/Salience network was driven by strong overlap with a single subnetwork, Ventral Attention B (**Figures 3C, A.3, and A.4**) at the level of individual participants. By contrast, the overlap of the pimfs-v was more variable across individuals and split among all three sub-networks (**Figures 3C, A.3, and A.4**). This observation was statistically supported by the pimfs-v having a larger Wasserstein distance (*W* = 6.07 × 10^6^, *p* < .001; pimfs-d: mean ± se = 0.0311 ± 0.0003, pimfs-v: mean ± se = 0.0325 ± 0.0003), which indicates decreased similarity and therefore greater variability between participants (**Materials and Methods**). As with the broad functional networks, the relationships between each component and the functional sub-networks did not differ based on whether or not the other component was present (*p*s > .37). The functional network dissociations between the pimfs components identified in individual participants also extended to meta-analytic fMRI data from over 10,000 neuroimaging experiments (Yeo et al., 2015); **Figure A.5**; **Materials and Methods**). Altogether, our analyses indicate that the pimfs-d and pimfs-v are functionally dissociable and have different connectivity fingerprints, despite being in close cortical proximity to one another (**Figure 1**).

### Components of the paraintermediate frontal sulcus can disappear on average surfaces: Implications for neuroimaging studies performing group analyses

The variable presence of the pimfs can affect neuroimaging studies aimed at assessing structural-functional correspondences using group analyses and averaged cortical surface reconstructions. For example, the putative “averaged” pimfs components are visible in the left, but not right, hemisphere of the commonly used fsaverage template (which is made from 39 participants, see https://surfer.nmr.mgh.harvard.edu/fswiki/FsAverage for additional details; **Figure 4A**). Notably,the fact that two components are visible in the left-hemisphere fsaverage template does not mean that the pimfs components are more common in the left hemisphere. Both in this adult sample (Willbrand, Jackson, et al., 2023) and a previous pediatric sample (Willbrand, Voorhies, et al., 2022), the incidence of pimfs-d and pimfs-v do not differ significantly across hemispheres.

**Figure 4.**
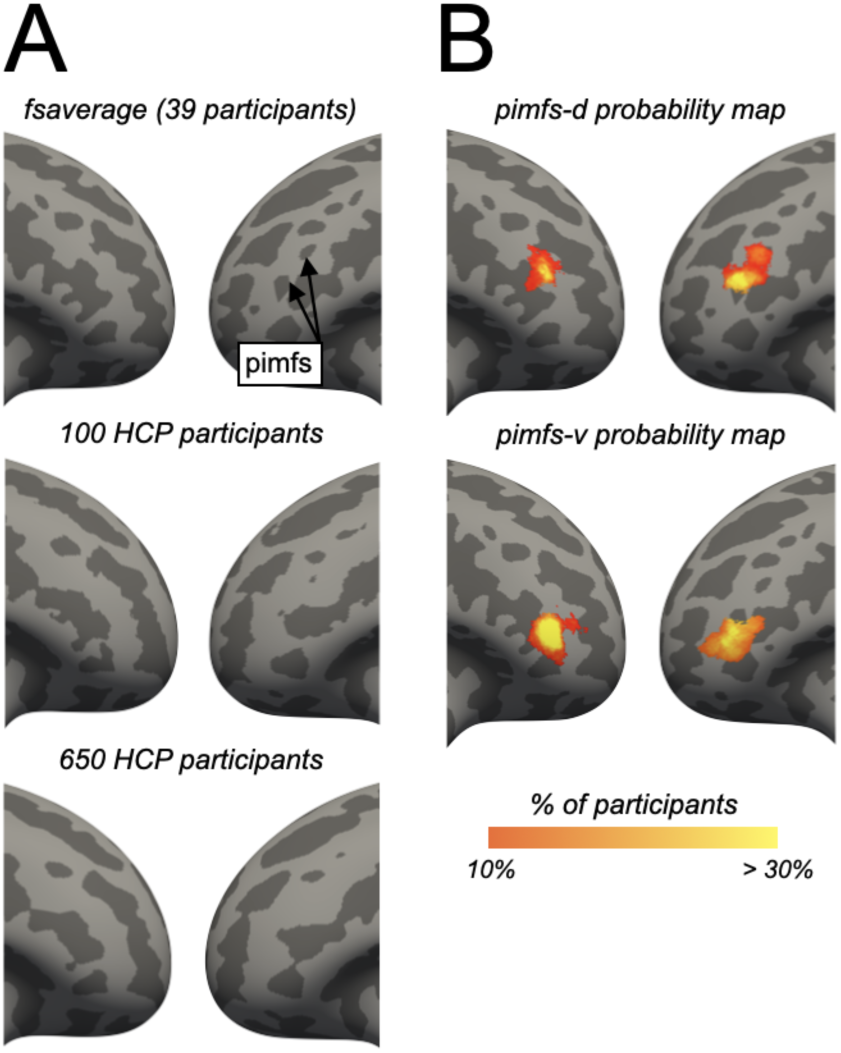
Paraintermediate frontal sulcal components can disappear on average surfaces. **A.** The variability of the pimfs in the aMFG can cause them to disappear when individual surfaces are averaged together. Surfaces are focused on LPFC, as in Figure 1. Top to bottom: i) fsaverage surface (39 participants), ii) 100 Human Connectome Project (HCP) participants, iii) 650 HCP participants. The disappearance of these sulci on average surfaces, which are often used for group analyses in neuroimaging research, emphasizes the importance of defining these structures on individual hemispheres (Figure 1). **B.** Probabilistic maximum probability maps (MPM; thresholded at 10% of vertex overlap across participants) of the pimfs-d (top) and pimfs-v (bottom) on the fsaverage surface (data from (Willbrand, Jackson, et al., 2023), showing that the likely location of the pimfs components do not necessarily align with clearly identifiable structures on average surfaces.

Beyond freely available templates such as the fsaverage surface, pimfs components can disappear when averaging randomly chosen cortical surfaces from large databases, such as the Human Connectome Project used in the present study (Glasser et al., 2013). For example, when randomly choosing either 100 or 650 HCP participants, the pimfs components are no longer visible in the left hemisphere (**Figure 4A**). This highlights the variability of the pimfs and, more generally, how anatomical variability could affect neuroimaging studies focused on anatomical-functional correspondences, thereby necessitating analyses at the level of individual participants.

## Discussion

### Overview

By applying a multimodal and multiscale approach based on classic criteria (Felleman & Van Essen, 1991; Kaas, 1997; Van Essen, 2003) to 249 pimfs labels from 72 participants, we demonstrated that the pimfs-d and pimfs-v—two variable sulci in anterior LPFC—are anatomically and functionally dissociable cortical structures (**Figure 1**). First, the pimfs-d and pimfs-v are morphologically dissociable based on their size (which is also the largest difference): the surface area of the pimfs-d is 15.73% smaller on average than the pimfs-v. Second, the pimfs-d and pimfs-v are dissociable based on architectural features (cortical thickness and myelination): the pimfs-d has a 2.48% higher thickness and 0.81% less myelination (T1-w/T2-w ratio proxy) on average than the pimfs-v. Third, the pimfs-d and pimfs-v are functionally dissociable based on data at the individual (Kong et al., 2019) and meta-analytic levels (Yeo et al., 2015).

The present study builds on the growing literature examining relationships between sulcal anatomy and the anatomical and functional organization of association cortices, including ventral temporal cortex (Weiner, 2019), posterior LPFC (Miller, Voorhies, et al., 2021), medial PFC (Amiez et al., 2013, 2021; Amiez & Petrides, 2014; Lopez-Persem et al., 2019), orbitofrontal cortex (Y. Li et al., 2015; Troiani et al., 2016, 2020), and posteromedial cortex (Willbrand, Parker, et al., 2022). In the sections below, we discuss these findings in the context of tertiary sulci serving as personalized coordinates for function and architecture in association cortices, and the impact of sulcal variability on regional anatomical and functional organization. We then discuss the limitations of this work and possible future directions.

### Tertiary sulci as “personalized coordinates” for function and architecture in association cortices

The present study shows that pimfs-d and pimfs-v are dissociable structures, which justifies the use of distinct anatomical labels for these sulci in current and future research. More generally, these results extend recent work proposing that tertiary sulci may serve as “personalized coordinates” for distinct functional and architectural areas in association cortices at the individual level (Miller, D’Esposito, et al., 2021). Specifically, with regard to the pimfs-d and pimfs-v in anterior LPFC, we find that these components represent a transition from attention-related networks to cognitive control-related networks (**Figure 3**).

To further contextualize the relationship between the pimfs components and anatomical/functional regions (albeit at the group level), we assessed whether probabilistic locations of the pimfs components also differed in their overlap with well-cited group-level modern multimodal parcellations derived based on measures such as topography, architecture, function, and connectivity (HCP-MMP; (Glasser et al., 2016) and a classic microstructural parcellation (Brodmann’s classic cytoarchitectonic parcellation (Brodmann, 1909; Van Essen, 2005); **Figure 5**; **Materials and Methods**). While we focus on the HCP-MMP and Brodmann cortical parcellations here, we show additional anatomically and/or functionally derived parcellations in **Figure A.6** (Fan et al., 2016; Foit et al., 2022; Scholtens et al., 2018; Vogt & Vogt, 1919; von Economo & Koskinas, 1925).

**Figure 5.**
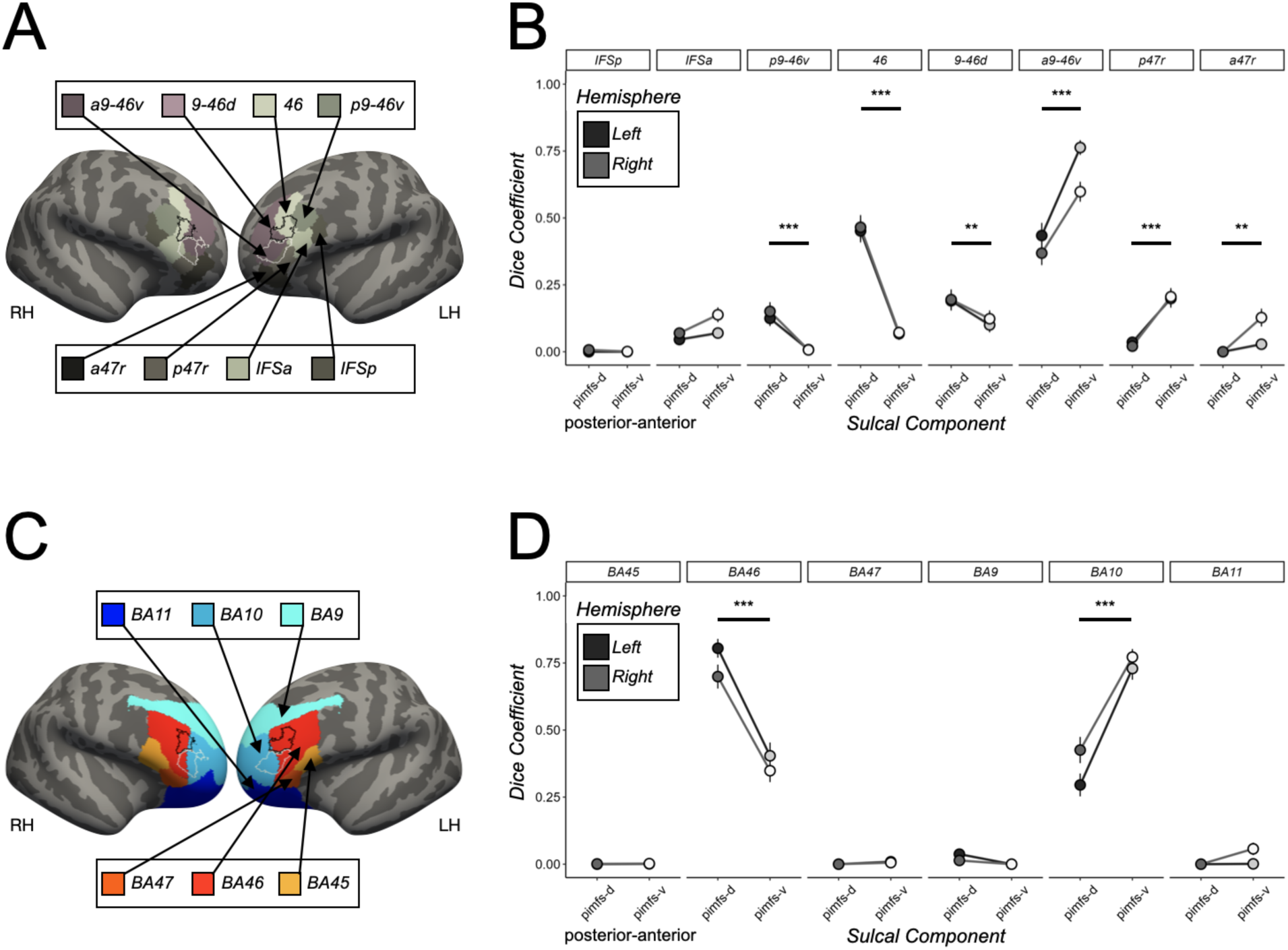
Pimfs components overlap with different regions in classic cytoarchitectonic and modern multimodal group-level cortical parcellations. **A.** Left (LH) and right (RH) fsaverage hemispheres displaying the relationship between the probabilistic location of the pimfs components (pimfs-d: black outline; pimfs-v: white outline; from (Willbrand, Jackson, et al., 2023) and eight LPFC regions in the HCP-MMP parcellation (Glasser et al., 2016). **B.** Dice coefficient overlap visualized as a function of sulcus (x-axis; pimfs-d: black, pimfs-v: white), HCP-MMP regions (subplots), and hemisphere (LH: darker shades; RH: lighter shades; see key). Large dots and error bars represent mean ± standard error (se). Horizontal lines and asterisks (*** *p* < .001, ** *p* < .01) indicate the significant post hoc pairwise comparisons from a sulcal component × region interaction [LME, predictors: sulcal component (pimfs-d and pimfs-v) × region × hemisphere (LH and RH); F(7, 1701) = 55.64, η2 = 0.19, *p* < .001]. This interaction was driven by the pimfs-d overlapping more with areas p9-46v (*d* = 0.72, *p* < .001), 9-46d (*d* = 0.30, *p* = .003), and 46 (*d* = 1.40, *p* < .001) and the pimfs-v overlapping more with areas a9-46v (*d* = 0.86, *p* < .001), p47r (*d* = 0.89, *p* < .001), and a47r (*d* = 0.55, *p* = .005). **C.** Same as A, except for the six LPFC regions in Brodmann’s cytoarchitectonic parcellation (Brodmann, 1909; Van Essen, 2005). **D.** Same format as B, but with Brodmann’s cytoarchitectonic parcellation. Again, there was a sulcal component × region interaction (F(5, 1215) = 93.19, η2 = 0.28, *p* < .001). This interaction was driven by the pimfs-d overlapping more with Brodmann area (BA) 46 (*d* = 1.12, *p* < .001; pimfs-d: mean ± se = 0.75 ± 0.03, pimfs-v: mean ± se = 0.38 ± 0.03) and pimfs-v overlapping more with BA 10 (*d* = 1.20, *p* < .001; pimfs-d: mean ± se = 0.36 ± 0.03, pimfs-v: mean ± se = 0.75 ± 0.03). Additional modern and classic cortical parcellations are shown in **Figure A.6**.

With regard to the HCP-MMP, pimfs-d showed similar overlap with area 46 (mean ± se = 0.46 ± 0.03) and area a9-46v (mean ± se = 0.40 ± 0.03), whereas the pimfs-v showed the highest overlap with area a9-46v (mean ± se = 0.68 ± 0.02; **Figure 5B**). In classic anatomical terms (Cunningham, 1892), these data suggest that the pimfs-d may serve as a limiting sulcus (that is, a boundary) separating areas 46 and a9-46v, while pimfs-v may serve as an axial sulcus (that is, co-localizing with) for area a9-46v (**Figure 5B**). On the other hand, overlap with Brodmann’s cytoarchitectural parcellation suggests that both pimfs components may be axial sulci for separate Brodmann areas (BAs). In this parcellation, the pimfs-d overlaps strongly with BA 46 (mean ± se = 0.75 ± 0.03), whereas the pimfs-v overlaps with BA 10 (mean ± se = 0.75 ± 0.03; **Figure 5D**). However, these results need further verification as these parcellations were derived at the group level (Glasser et al., 2016; Van Essen, 2005) and their identification was observer-dependent (Brodmann, 1909; Glasser et al., 2016). Future work with post-mortem data, along with individual-level, observer-independent MRI data (e.g., (Amunts et al., 2020) is needed to further investigate this relationship.

The putative overlap of pimfs-v with parcels in the HCP-MMP and Brodmann atlases links this sulcus to the fMRI literature on reasoning. The HCP-MMP parcel with which pimfs-v overlapped most strongly (area a9-46v) was shown to be functionally dissociable from nearby areas by Glasser and colleagues on the basis of activation during the performance of a relational reasoning task (the relational-match contrast; (Glasser et al., 2016). Similarly, the Brodmann area with which pimfs-v putatively overlaps (BA 10) has been routinely reported for a variety of reasoning tasks, especially in relation to rostrolateral PFC (RLPFC), a functionally identified region implicated in reasoning (Christoff & Gabrieli, 2000; Holyoak & Monti, 2021; Koechlin et al., 1999; Ramnani & Owen, 2004; Smith et al., 2007; Urbanski et al., 2016; Vendetti & Bunge, 2014; Wendelken et al., 2008; Westphal et al., 2016, 2019). This correspondence suggests that pimfs-v may lie within an area of LPFC functionally related to reasoning (i.e., RLPFC). Future work delineating sulcal anatomy and task-related fMRI activation at the individual level is needed to confirm this overlap.

In sum, the pimfs components may be axial or limiting sulci, depending on the parcellation (**Figures 4, 5, and A.6**). These results, alongside prior work (Y. Li et al., 2015; Lopez-Persem et al., 2019; Miller, D’Esposito, et al., 2021; Miller, Voorhies, et al., 2021; Troiani et al., 2016, 2020; Weiner, 2019; Willbrand, Parker, et al., 2022), emphasize that future parcellations should incorporate individual-level sulcal definitions to more accurately delineate regions. Further, the pimfs components overlap with HCP-MMP areas called a “hotspot of individual variability” in terms of topography, architecture, function, and connectivity (Glasser et al., 2016). Therefore, a goal for future research is to investigate how individual-level sulcal variability in LPFC relates to individual-level variability in these metrics and parcellations, especially given that two pimfs components are not present in every hemisphere.

### The impact of sulcal variability on regional anatomical and functional organization

Prior work has demonstrated that the presence or absence of sulci in association cortices impacts the location of cytoarchitectural areas, task-related activation, and functional networks, particularly in medial PFC (Amiez et al., 2013, 2021; Amiez & Petrides, 2014; Lopez-Persem et al., 2019). Although the present investigation focused primarily on the pimfs-d and pimfs-v in cases where these structures were present, many but not all individuals have both pimfs components in a given hemisphere, as noted above (Willbrand, Jackson, et al., 2023; Willbrand, Voorhies, et al., 2022). This variability also has behavior implications: our previous work indicated that the presence of the left pimfs-v is associated with 21-34% better reasoning scores in pediatric and adult samples (Willbrand, Jackson, et al., 2023; Willbrand, Voorhies, et al., 2022).

Prior fMRI research on reasoning has most consistently emphasized the role of left RLPFC in reasoning (Assem et al., 2020; Bunge et al., 2009; Christoff et al., 2001; Green et al., 2006, 2010; Hartogsveld et al., 2018; Hobeika et al., 2016; Urbanski et al., 2016; Wendelken et al., 2017). Given our prior work revealing better reasoning performance overall among participants with a left, but not right, pimfs-v (Willbrand, Jackson, et al., 2023; Willbrand, Voorhies, et al., 2022), future work should assess whether the incidence of this sulcal component relates to activation of RLPFC—and whether this mediates the behavioral difference seen between individuals who do or do not possess a left pimfs-v component. Further, once improved cytoarchitectural definitions fully characterize the aMFG (Amunts et al., 2020; Bludau et al., 2014; Bruno et al., 2022; Wojtasik et al., 2020), future investigations could relate the presence/absence of the pimfs components to shifts in cytoarchitectonic regions (**Figure 5D**), given previous findings for sulcal variations in medial PFC (Amiez et al., 2021).

### Limitations

Although the present work contained a relatively large sampling of sulci for individual-level analyses (the identification of 249 pimfs informed by the location of additional LPFC sulci, resulting in 2,985 sulci defined), the primary limitation was the sample size (72 participants; 144 hemispheres). The sample sizes of studies involving manually defined sulci in individual participants (e.g., (Amiez et al., 2006, 2018, 2019; Borst et al., 2016; Cachia et al., 2014; Garrison et al., 2015; Hopkins et al., 2021; Lopez-Persem et al., 2019; Miller, Voorhies, et al., 2021; Nakamura et al., 2020; Voorhies et al., 2021; Weiner et al., 2014; Willbrand, Parker, et al., 2022; Willbrand, Voorhies, et al., 2022; Yao et al., 2022; Zlatkina et al., 2016) are limited by the time investment and anatomical expertise required to label them. With the advent of improved methods to automatically define sulci (e.g., (Borne et al., 2020; Lyu et al., 2021; Willbrand, Parker, et al., 2022), such sulcal-based studies can begin to increase their scope and scale. However, these methods are still developing, given the large (and still growing) number of sulci identifiable in the human (and non-human hominoid) cerebral cortex and the uniqueness of sulcal patterning at the individual level. Therefore, for the time being, manual and automatic methods must be used in tandem to further delineate the complex sulcal patterning of the cortex.

### Future directions

Altogether, the present findings and our prior work demonstrate the feasibility of applying this multimodal approach for dissociating sulci from one another and also determining the relevance of these structures across association cortices (Miller, Voorhies, et al., 2021; Willbrand, Parker, et al., 2022). Future work should test these and additional methodologies in other regions and samples to determine the generalizability of these relationships and explore new questions. For example, although sulci appear in gestation (Chi et al., 1977), sulcal morphology does change during child development (Alemán-Gómez et al., 2013; Klein et al., 2014; Meng et al., 2014; Raznahan et al., 2011; Vandekar et al., 2015; Willbrand, Ferrer, et al., 2023), Thus, the differences in thickness, myelination, and surface area of the pimfs components may relate to underlying differential rates of development. Additionally, we identified a small difference in the myelination between the pimfs components (via the T1-w/T2-w ratio proxy), but it is an open question as to whether there are differences between the pimfs components in terms of white matter projections that could explain or contribute to differences in functional connectivity profiles. Exploring these multiple possibilities could provide insight into how LPFC hierarchies develop on a microscale. Further, these relationships between the pimfs components may change with age: that is, do the pimfs components begin as morphologically, anatomically, and functionally distinct at birth, or do they differentiate during infant/child development? These anatomical and functional relationships may also differ as a function of psychiatric or neurological conditions that have roots in prenatal development when the sulci first form (Cachia et al., 2021; Chi et al., 1977). Finally, given the variability of the pimfs and their unique location in LPFC at the convergence between dorsal-ventral and rostral-caudal axes in LPFC, these sulci may serve as convergence zone for anatomical and functional gradients in LPFC (Badre & D’Esposito, 2009; Miller, D’Esposito, et al., 2021; Miller, Voorhies, et al., 2021; Nee & D’Esposito, 2016).

### Conclusion

To conclude, the present study further supports the claim that sulci can serve as a powerful tool, providing personalized anatomical coordinates (Miller, D’Esposito, et al., 2021) that “precision neuroimaging” studies (Gratton et al., 2022) can leverage to improve understanding of neuroanatomical-functional relationships at the individual level.

## Appendix

**Figure A.1.**
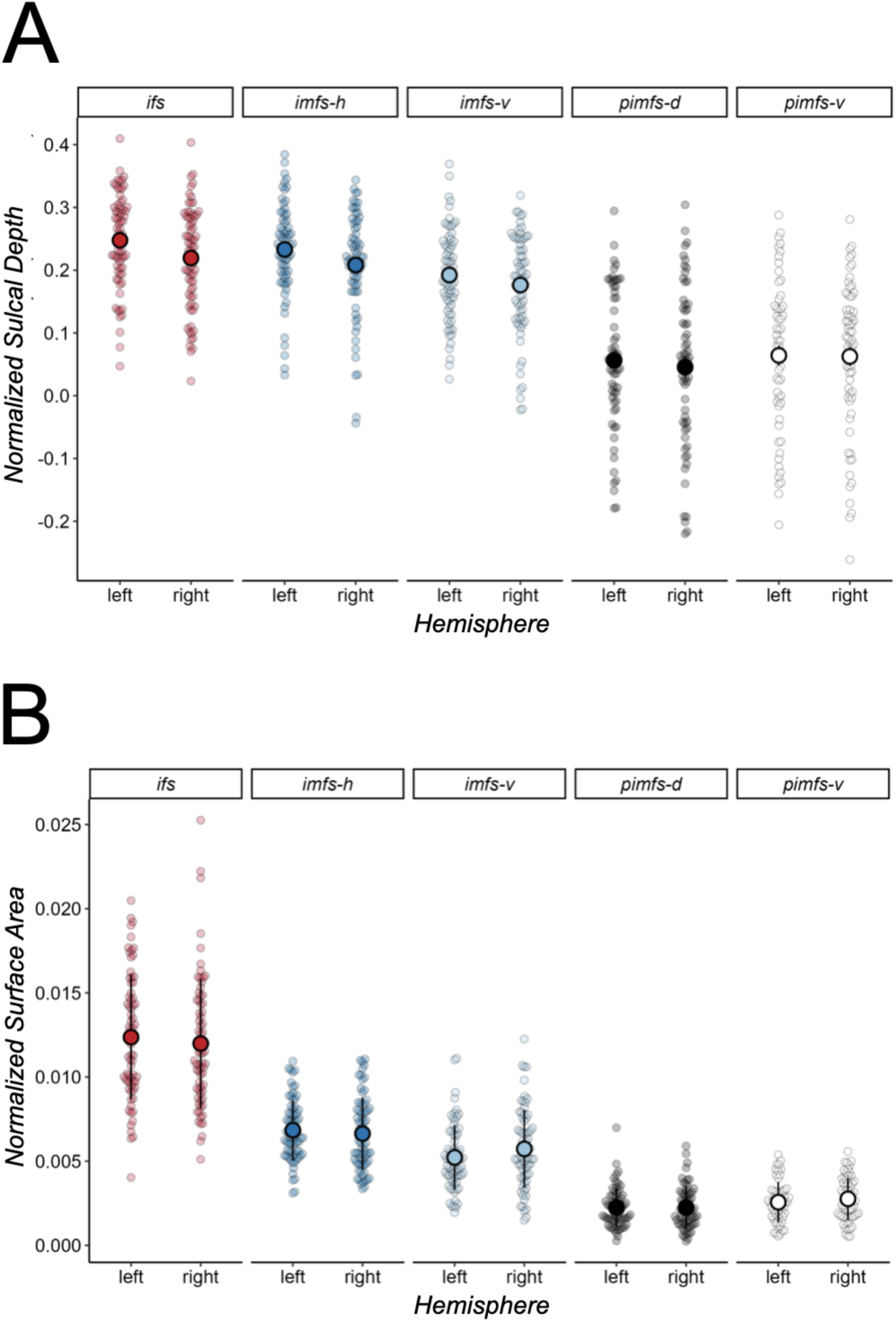
Sulcal depth and surface area of pimfs in relation to surrounding anterior LPFC sulci. **A.** Mean and standard deviation (large dot and bar) of the normalized depth of the two pimfs components and three more prominent sulci surrounding them (ifs, imfs-h, imfs-v; facets) in each hemisphere (x-axis). Depth is normalized to the maximum depth in each hemisphere (Materials and Methods). Individual dots represent individual values for each participant. Sulci are colored according to Figure 1A. **B.** Same as A, but for surface area (normalized by cortical surface area; Materials and Methods). The pimfs components are far shallower and smaller than the sulci surrounding them (Amiez et al., 2019; Lopez-Persem et al., 2019; Miller, D’Esposito, et al., 2021; Miller et al., 2020; Miller, Voorhies, et al., 2021; Sanides, 1964; Voorhies et al., 2021; Weiner, 2019; Weiner et al., 2014; Welker, 1990; Willbrand, Parker, et al., 2022; Yao et al., 2022).

**Figure A.2.**
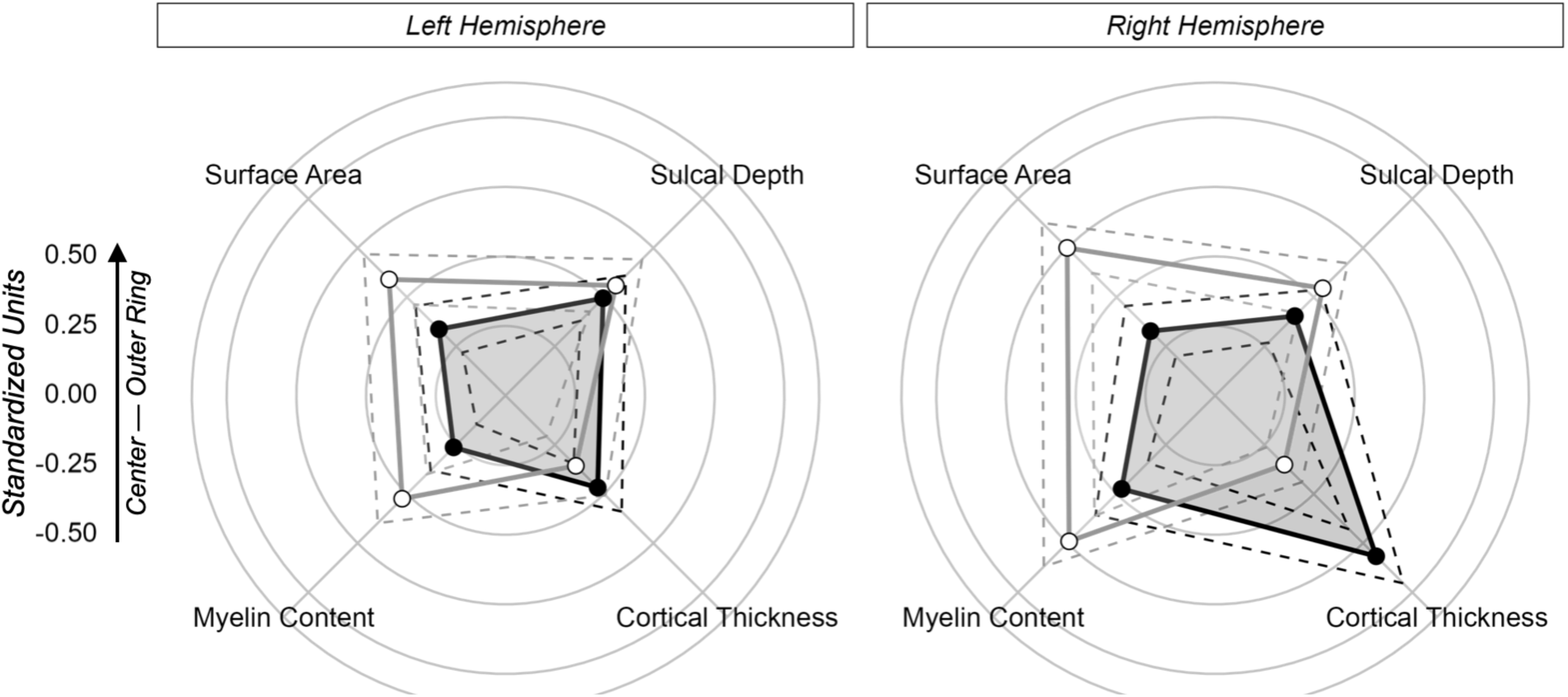
Morphological and architectural features of the pimfs components in each hemisphere separately. Polar plot showing the mean morphological (top features) and architectural (bottom features) values for the pimfs components in the left (left facet) and right (right facet) hemispheres separately. Solid lines and dots represent the means. Dashed lines represent ± standard error. Lines and dots are colored by sulcal component (pimfs-d: black, pimfs-v: white/gray). Each concentric circle corresponds to the units shown to the left, which are standardized to allow for these metrics to be plotted together.

**Figure A.3.**
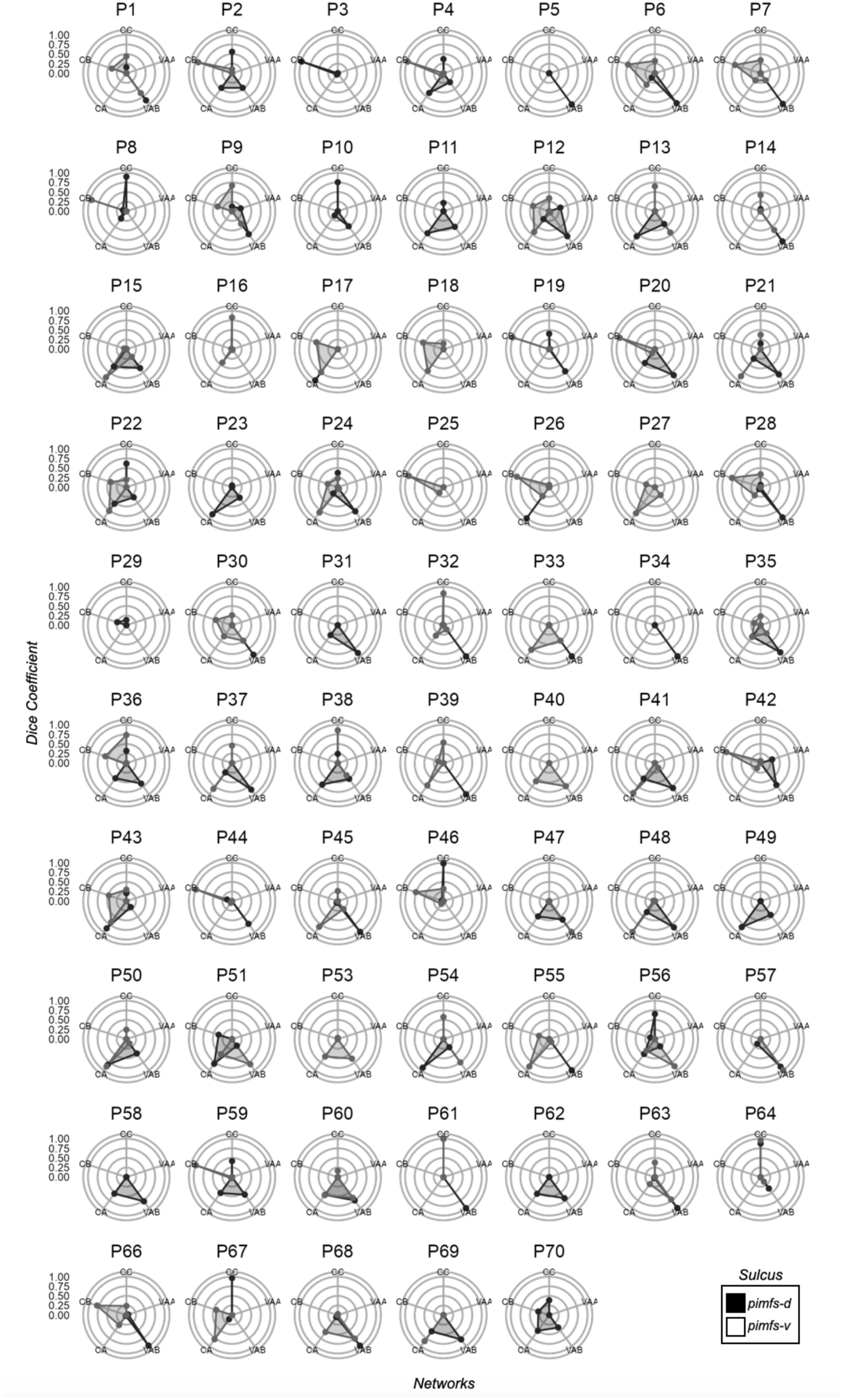
Functional connectivity fingerprints of pimfs components in individual left hemispheres. Polar plots showing the connectivity fingerprints of the pimfs-d and pimfs-v with the Control (C) and Ventral Attention (VA) sub-networks in the left hemisphere of all participants with at least one pimfs component (N = 68). The closer to the periphery of the circle, the higher the Dice coefficient (numbers on the left correspond to the Dice coefficient value at each concentric circle).

**Figure A.4.**
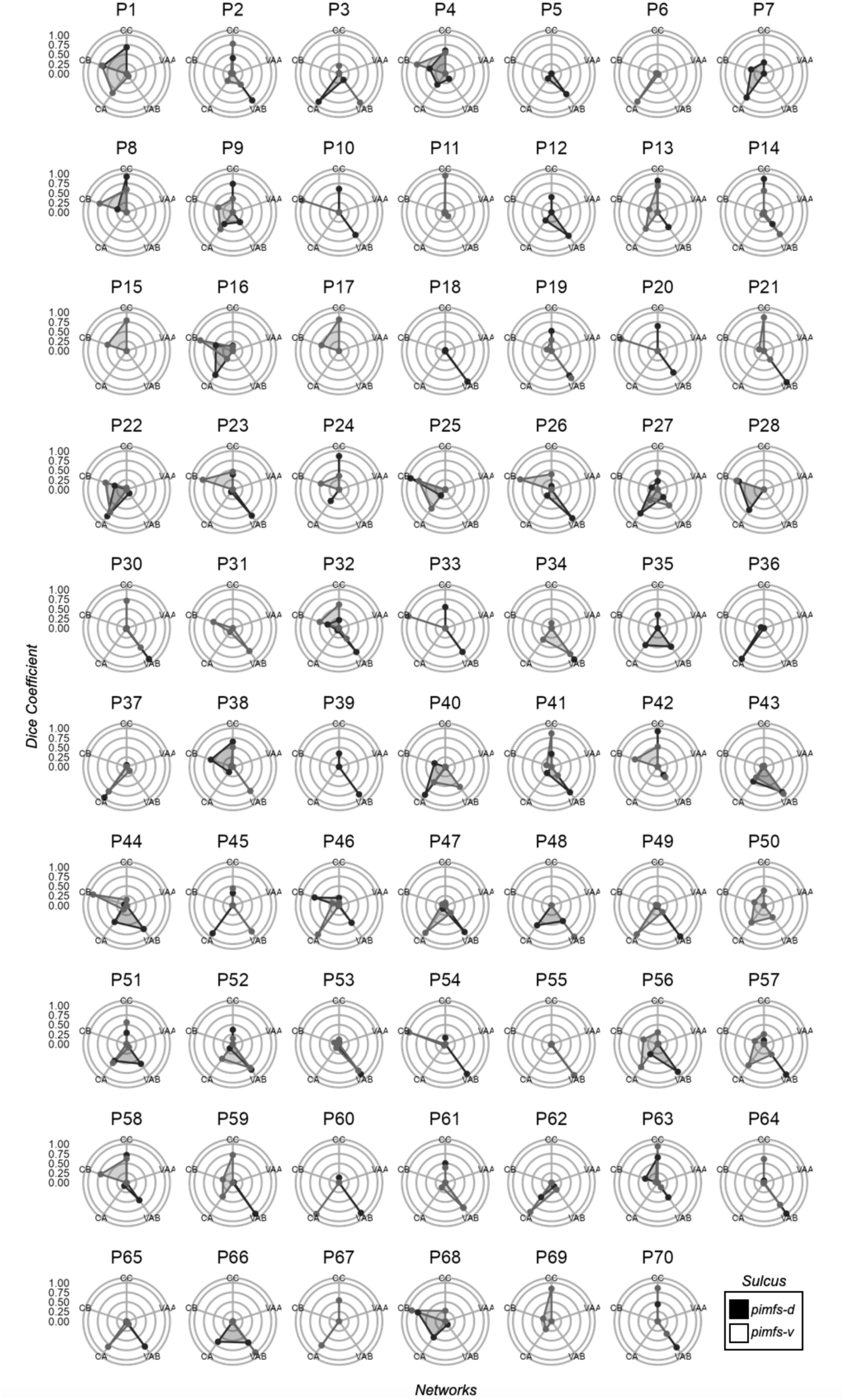
Functional connectivity fingerprints of pimfs components in individual right hemispheres. Polar plots showing the connectivity fingerprints of the pimfs-d and pimfs-v with the Control (C) and Ventral Attention (VA) sub-networks in the right hemisphere of all participants with at least one pimfs component (N = 69). The closer to the periphery of the circle, the higher the Dice coefficient (numbers on the left correspond to the Dice coefficient value at each concentric circle).

**Figure A.5.**
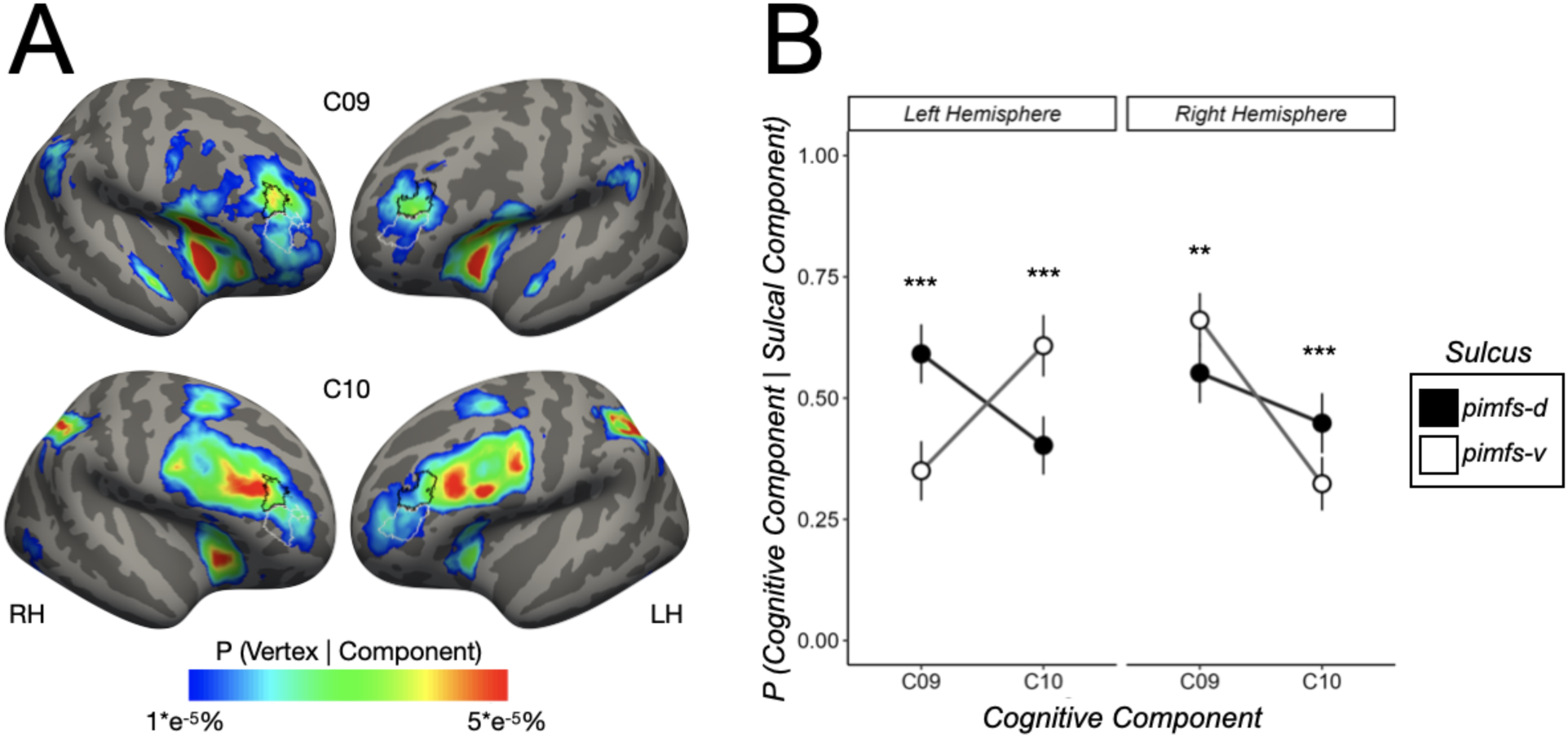
Cognitive component profiles of pimfs components differ. Using an expectation-maximization algorithm (Materials and Methods), we quantified the posterior probability between each of the 14 cognitive components (from a meta-analysis of fMRI experimental tasks; (Yeo et al., 2015) being associated with each pimfs component from each hemisphere of each individual (all in fsaverage space). In A and B, we only show the two cognitive components that displayed significant differences between the pimfs cognitive components: Component 9 (C09; inhibitory control) and Component 10 (C10; executive function). **A.** Left (LH) and right (RH) fsaverage hemispheres displaying the relationship between the probabilistic location of the pimfs components (pimfs-d: black outline; pimfs-v: white outline; from (Willbrand, Jackson, et al., 2023) and the two cognitive components (heatmaps) that displayed significant differences between the pimfs components. **B.** Posterior probability (P) visualized as a function of cognitive component (x-axis), sulcal component (pimfs-d: black; pimfs-v: white), and hemisphere (left hemisphere: left facet; right hemisphere: right facet). Large dots and error bars represent the means ± standard errors. These mean dots are connected by lines to help indicate the sulcal component × cognitive component × hemisphere interaction from an LME [predictors: sulcal component (pimfs-d and pimfs-v) × 14 cognitive components × hemisphere (LH and RH); F(13, 3159) = 8.51, η2 = 0.03, *p* < .001]. Asterisks (** *p* < .01; *\*\*\* p* < .001) indicate the significance of post hoc pairwise comparisons on the sulcal component × cognitive component × hemisphere interaction. This interaction was driven by a hemispheric dissociation in cognitive component and sulcal component probability. In the left hemisphere, pimfs-d loaded more onto C09 (*d* = 0.51, *p* < .001) while pimfs-v loaded more onto C10 (*d* = 0.43, *p <* .001). In the right hemisphere, pimfs-d loaded more onto C10 (*d =* 0.27, *p* < .001) while pimfs-v loaded more onto C09 (*d* = 0.23, *p =* .001).

**Figure A.6.**
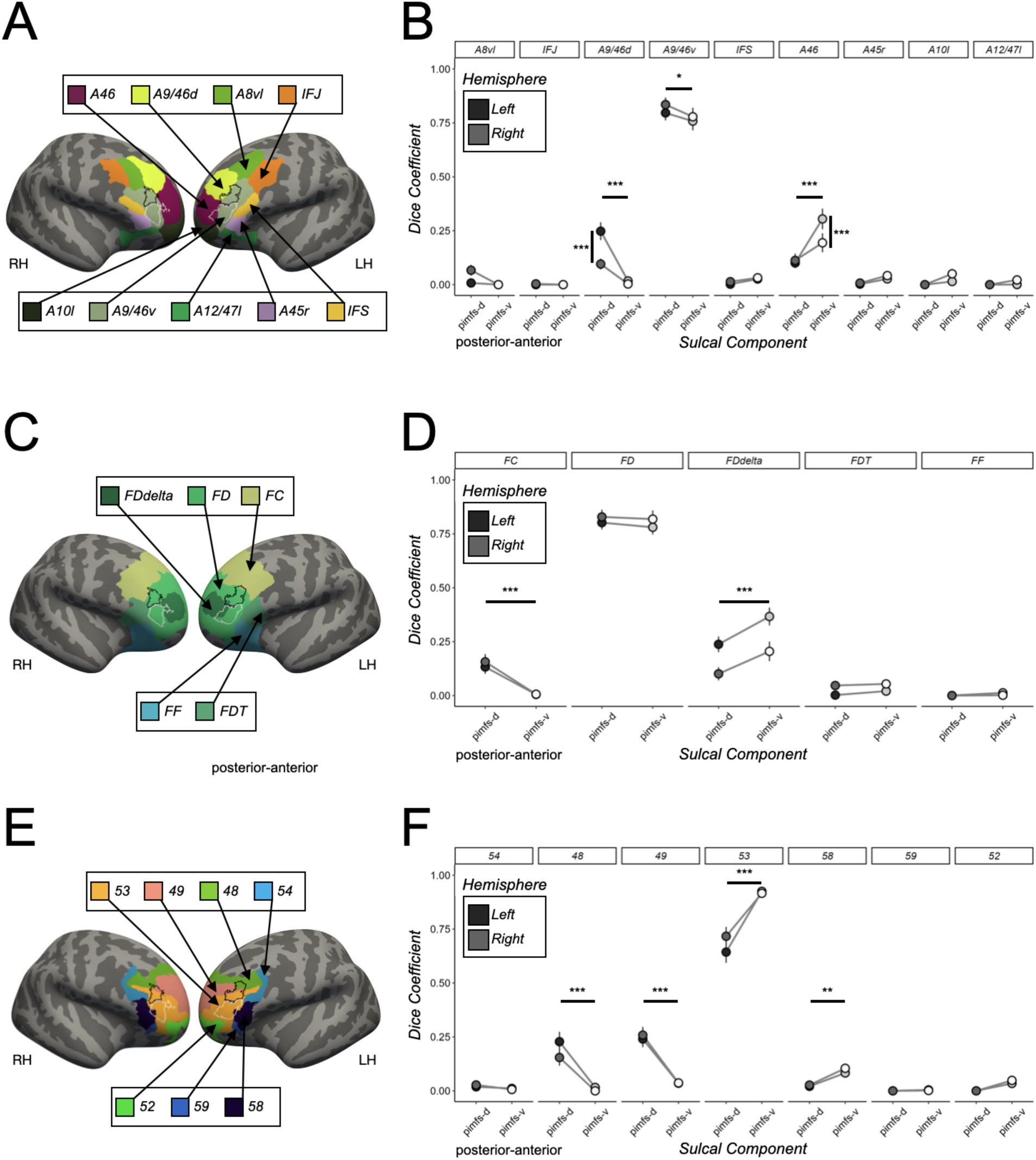
Pimfs components in relation to different areas in additional cortical parcellations. **A.** Left (LH) and right (RH) hemisphere fsaverage surfaces displaying the relationship between the probabilistic location of the pimfs components (pimfs-d: black outline; pimfs-v: white outline; from (Willbrand, Jackson, et al., 2023) and nine LPFC regions in the Brainnetome resting-state functional connectivity-based parcellation (Fan et al., 2016). **B.** Dice coefficient overlap visualized as a function of sulcus (x-axis; pimfs-d: black, pimfs-v: white), Brainnetome regions (subplots), and hemisphere (LH: darker shades; RH: lighter shades; see key). Large dots and error bars represent mean ± standard error (se). Horizontal lines and asterisks (*** *p* < .001, ** *p* < .01, * *p* < .05) indicate the significant post hoc pairwise comparisons from the sulcal component × region interaction [LME: predictors: sulcal component (pimfs-d and pimfs-v) × region × hemisphere (LH and RH); F(8, 1944) = 14.49, η2 = 0.06, *p* < .0001]. This interaction was driven by the pimfs-d overlapping more with areas A8vl, A9/46d, and A9/46v (*d*s > 0.15, *p*s < .028) and the pimfs-v overlapping more with area 46 (*d* = 0.47, *p* < .001). Vertical lines and asterisks indicate the significant post hoc pairwise comparisons from the sulcal component × region × hemisphere interaction (F(8, 1944) = 2.76, η2 = 0.01, *p* < .005). **C.** Same as A, except for the five LPFC regions in Von Economo and Koskinas’ cytoarchitectonic parcellation (Scholtens et al., 2018; von Economo & Koskinas, 1925). **D.** Same format as B, but with Von Economo and Koskinas’ cytoarchitectonic parcellation. Again, there was a sulcal component × region interaction (F(4, 972) = 11.88, η2 = 0.05, *p* < .001). This interaction was driven by the pimfs-d overlapping more with area FC (*d* = 0.73, *p* < .001) and the pimfs-v overlapping more with area FDdelta (*d* = 0.36, *p* < .001). **E.** Same as A, except for the seven LPFC regions in Vogt and Vogt’s myeloarchitectonic parcellation (Foit et al., 2022; Vogt & Vogt, 1919). **F.** Same format as B, but with Vogt and Vogt’s myeloarchitectonic parcellation. Again, there was a sulcal component × region interaction (F(6, 1458) = 45.42, η2 = 0.06, *p* < .001). This interaction was driven by the pimfs-d overlapping more with areas 48 and 49 (*d*s > 0.30, *p*s < .001) and the pimfs-v overlapping more with areas 53 and 58 (*d*s > 0.52, *p*s < .003).

## Data availability statement

The processed data required to perform all statistical analyses and reproduce all figures used for this project will be made freely available on GitHub upon publication (https://github.com/cnl-berkeley/stable_projects). The analysis pipelines used for this project are available on Open Science Framework (https://osf.io/7fwqk/). Anonymized neuroimaging data for the HCP participants are available at ConnectomeDB (db.humanconnectome.org). Requests for any additional information should be directed to the Corresponding Author, Kevin Weiner.

## Acknowledgments

We thank Jacob Miller, Willa Voorhies, Jewelia Yao, Samantha Jackson, and Szeshuen Chen for their prior assistance in defining LPFC sulci. We also thank Jacob Miller for helping develop the analysis pipelines implemented in the present work.

